## Supplementary material for "MAP1B Regulates Cortical Neuron Interstitial Axon Branching Through the Tubulin Tyrosination Cycle": Source Data

### Source Data 1

#### A Complete data for Figure 1B

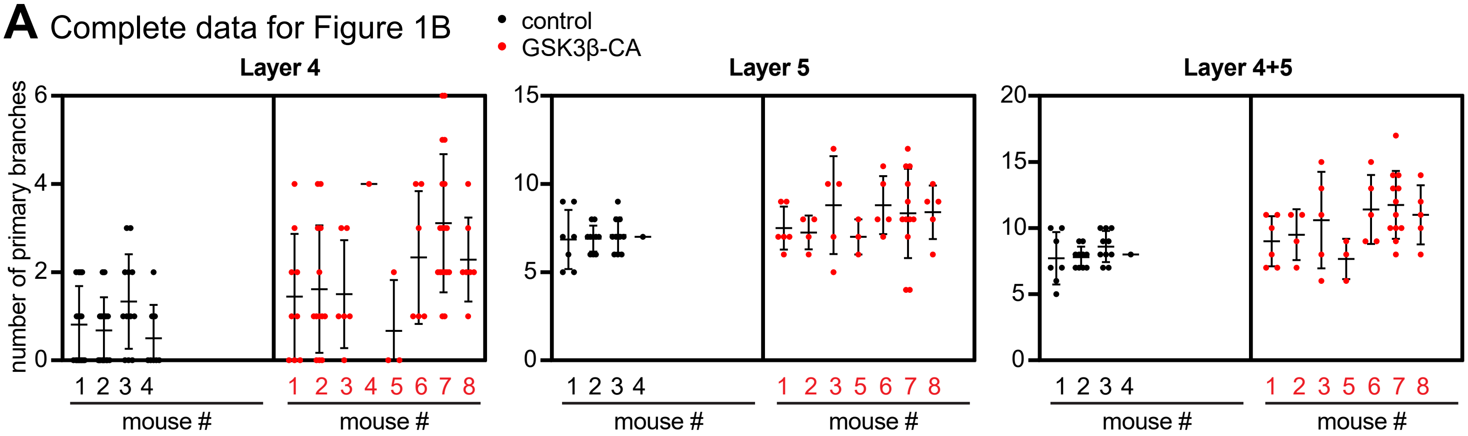


#### B Complete data for Figure 1C

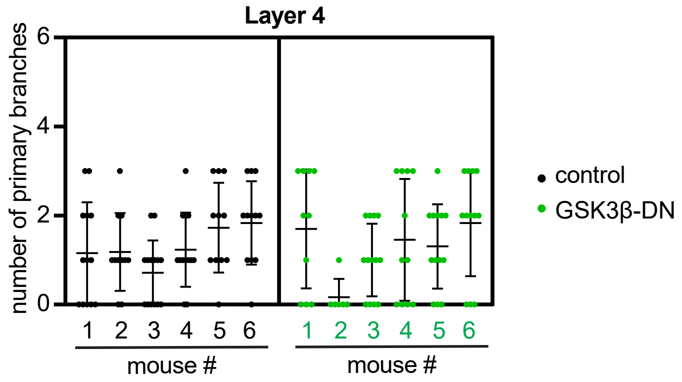


#### C Complete data for Figure 1E & Figure EV1F

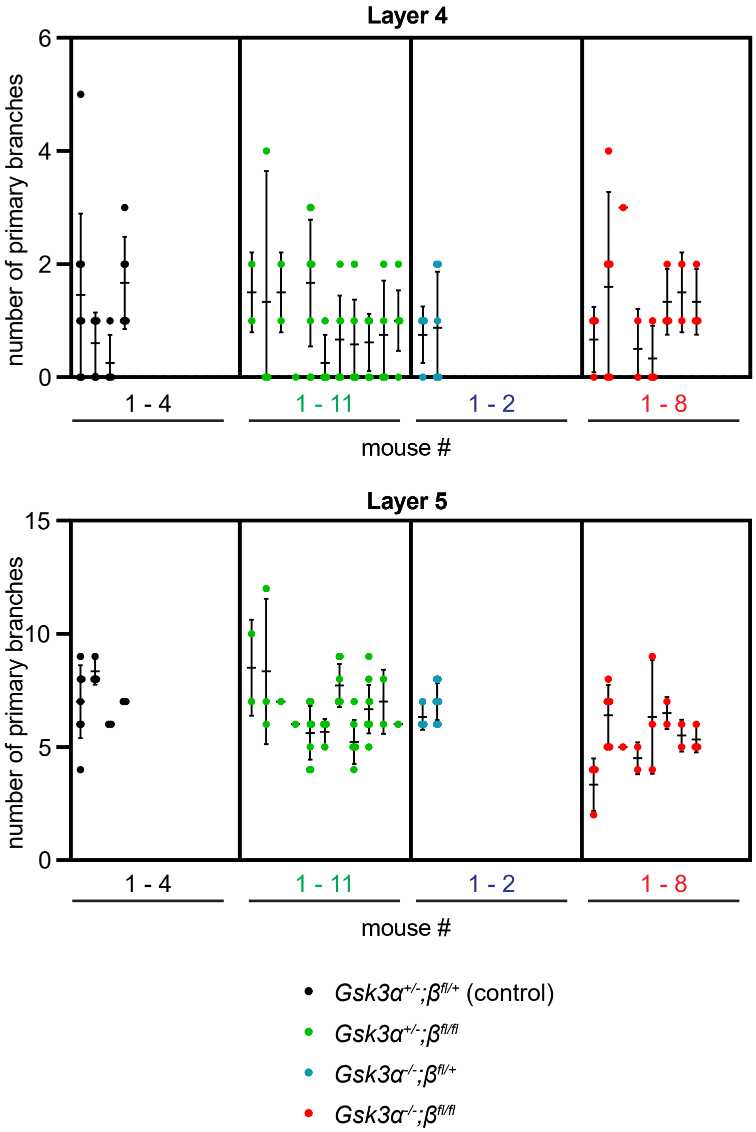


#### D Complete data for Figure EV1B

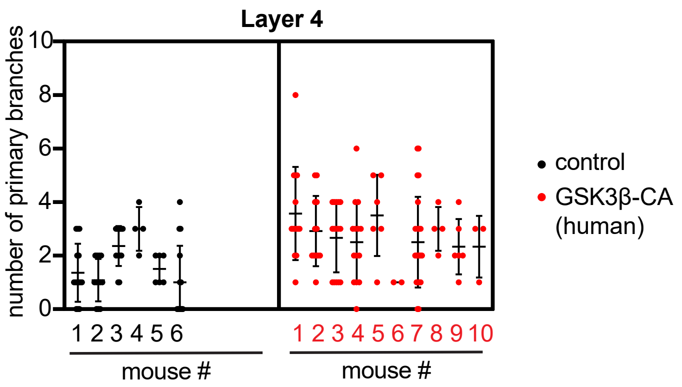


#### E Complete data for Figure EV1D

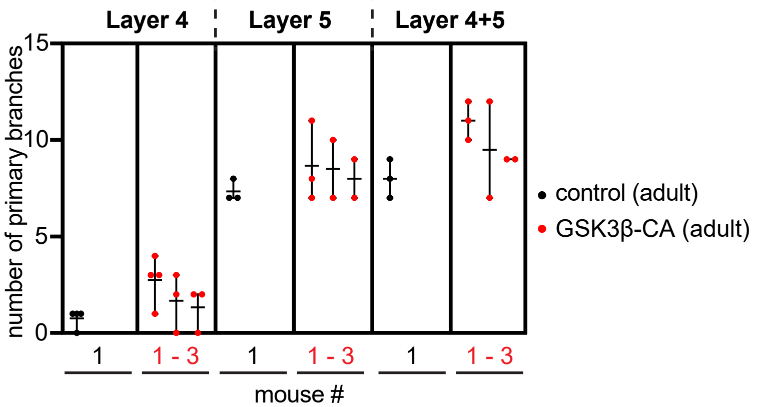

### Source Data 2

#### A Complete data for Figure 2A

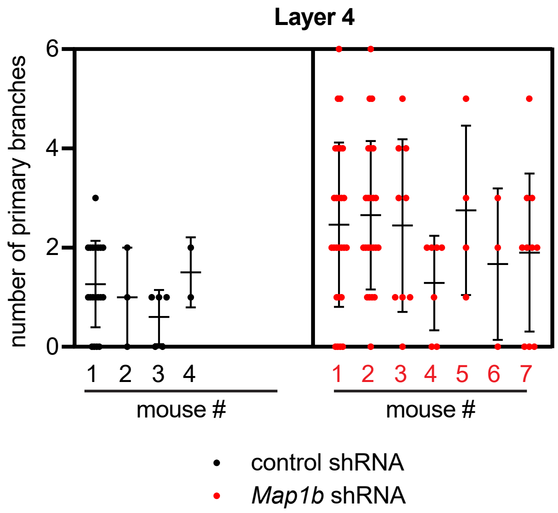


#### B Complete data for Figure 2C

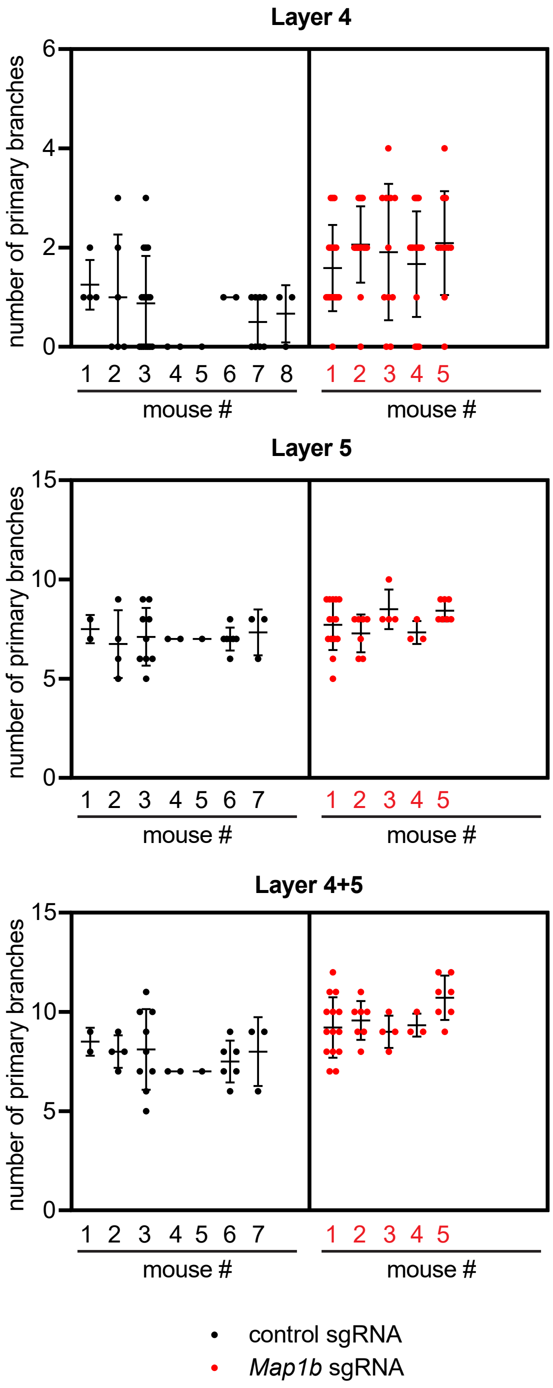


#### C Complete data for Figure 2E

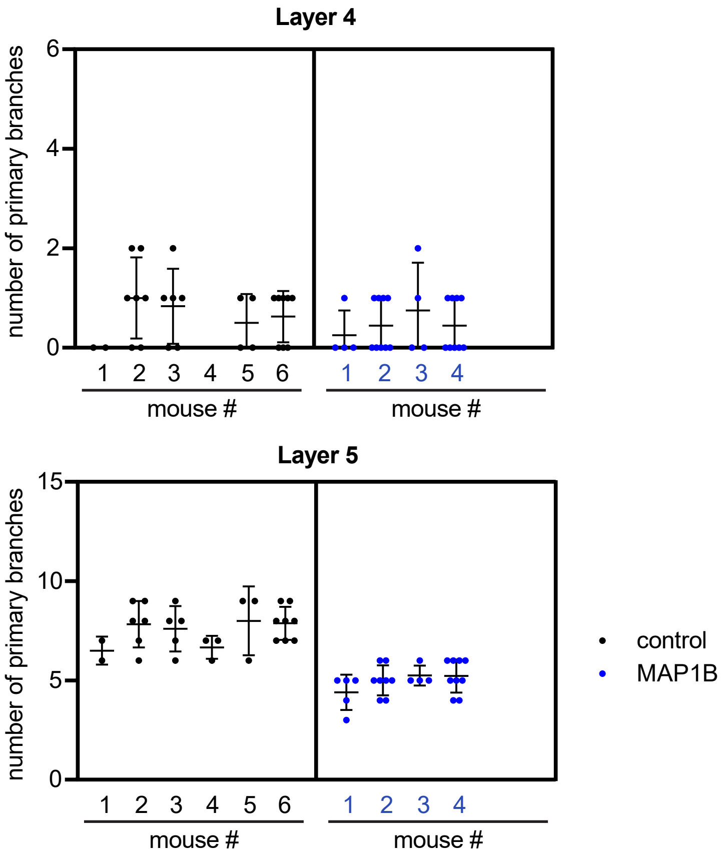

### Source Data 3

#### A Complete data for Figure 3A

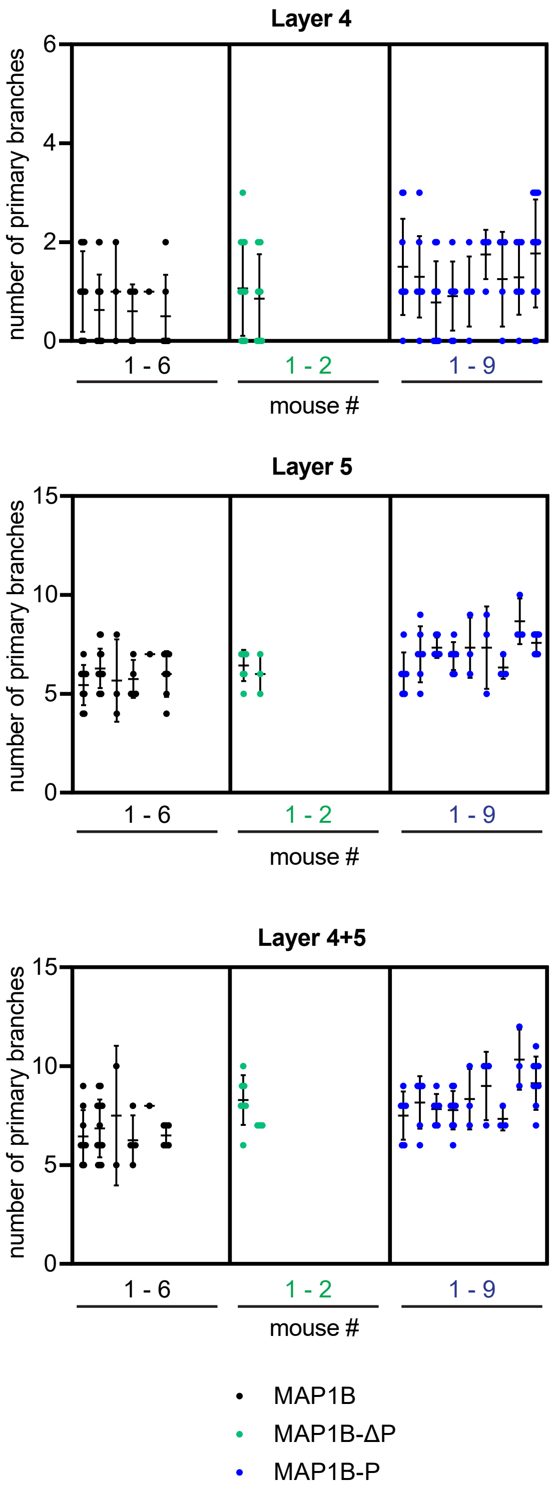


#### B Complete data for Figure 3C

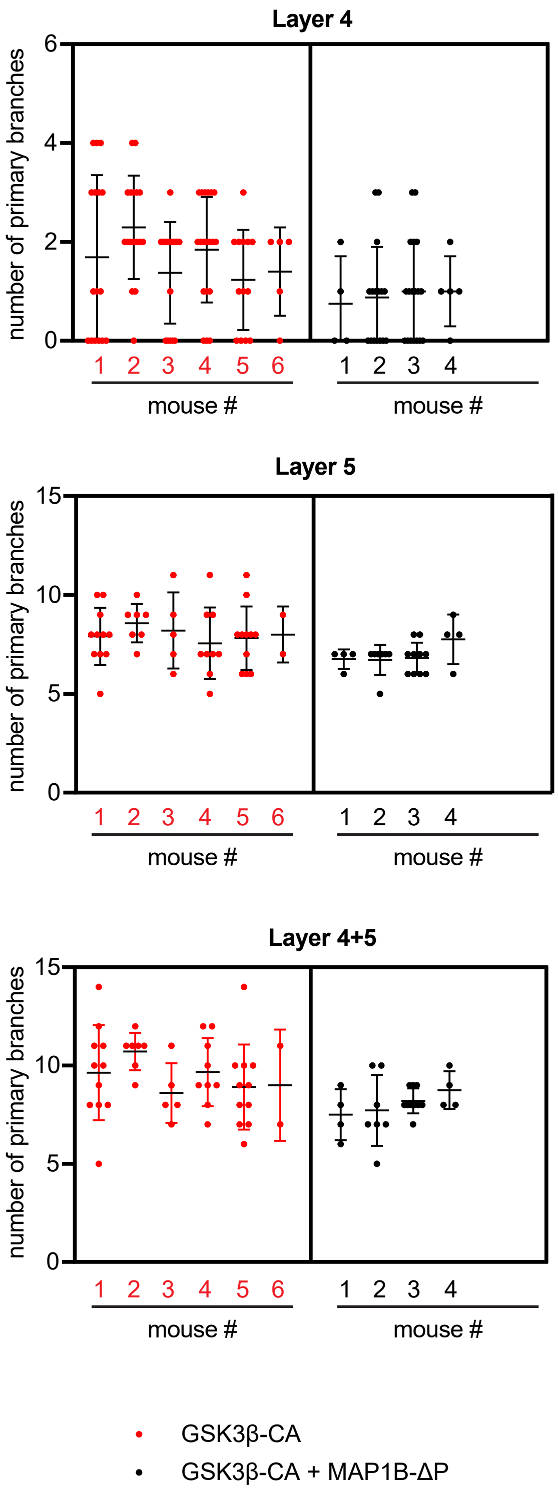

### Source Data 4

#### A Complete data for Figure 4C

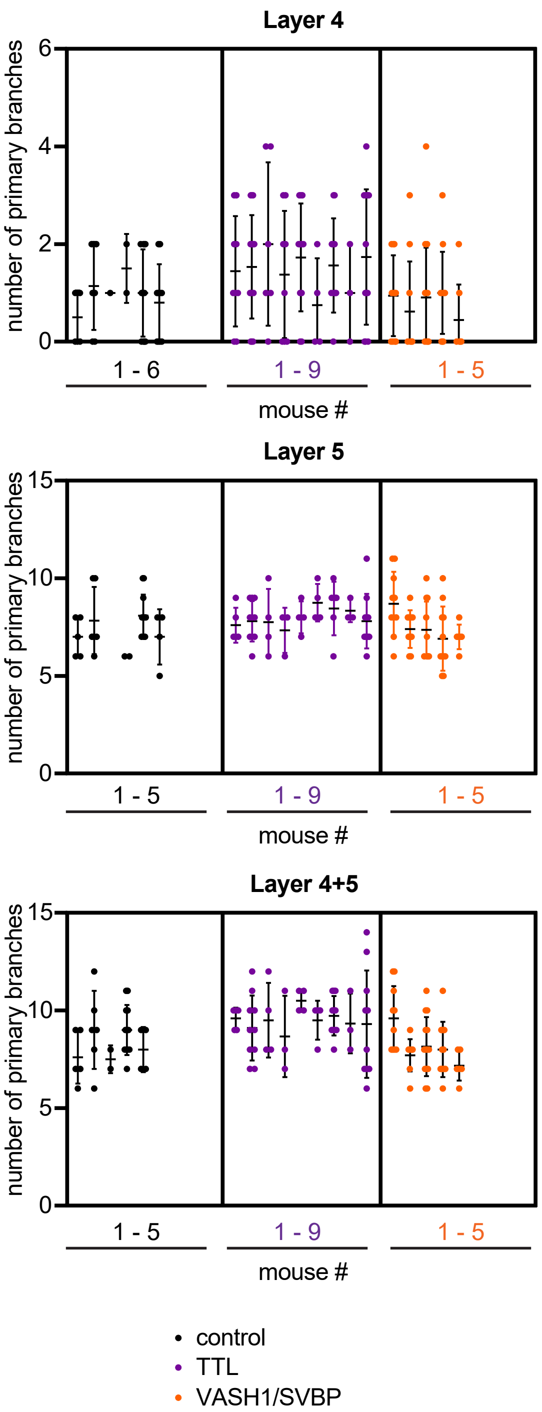


#### B Complete data for Figure 4F

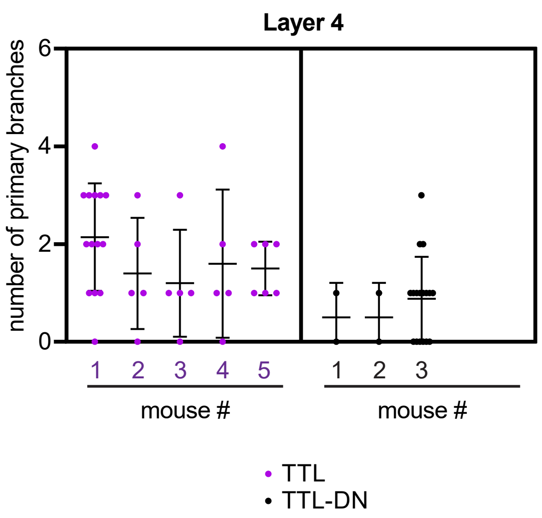

### Source Data 5

#### A Complete data for Figure 5A

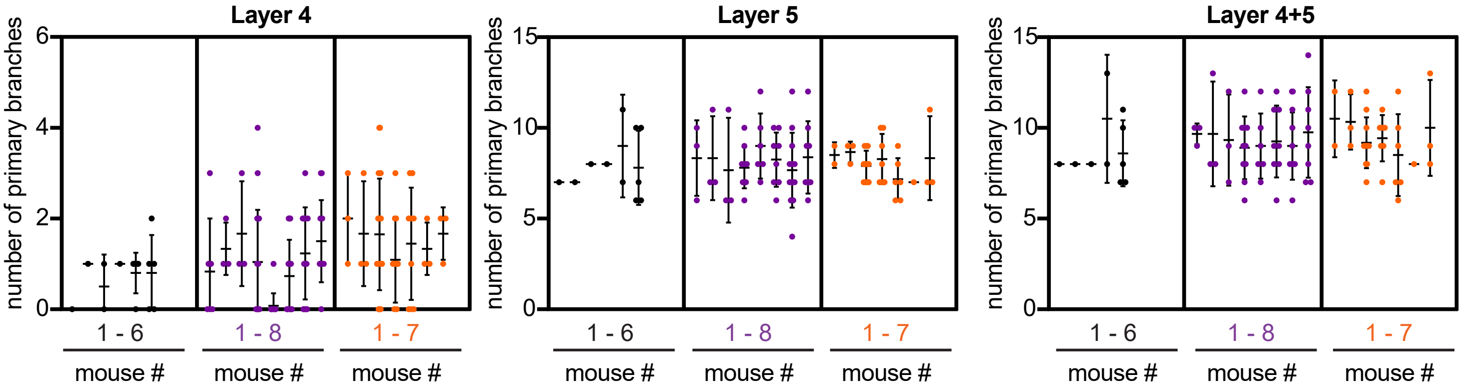


#### B Complete data for Figure 5C

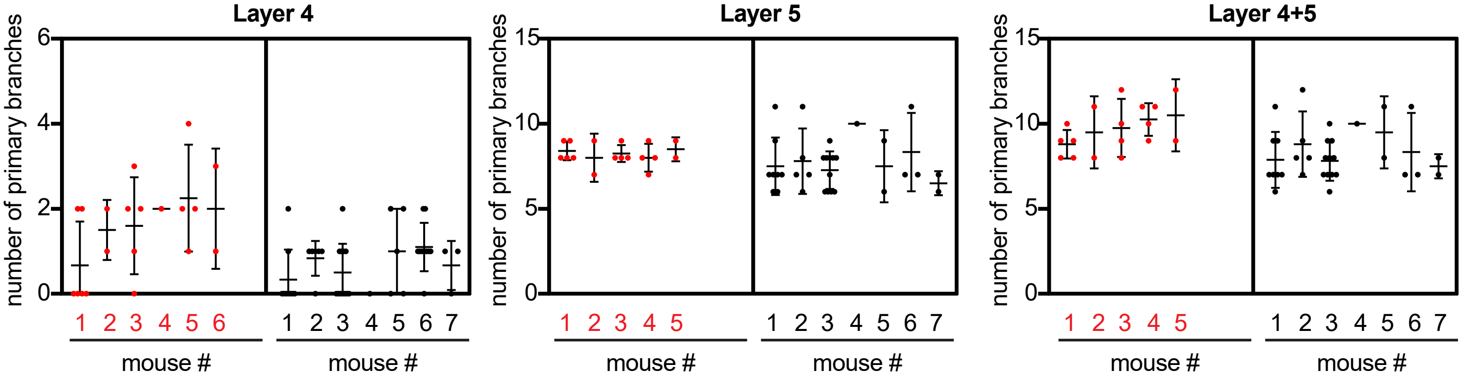


#### C Complete data for Figure EV5A

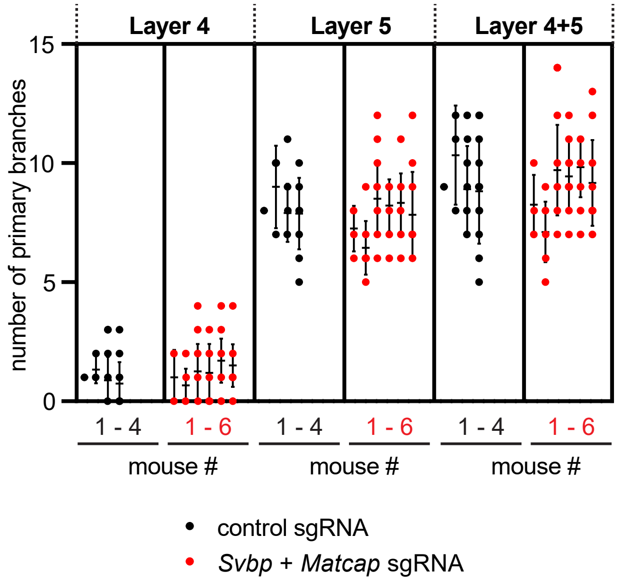


#### D Complete data for Figure 6A

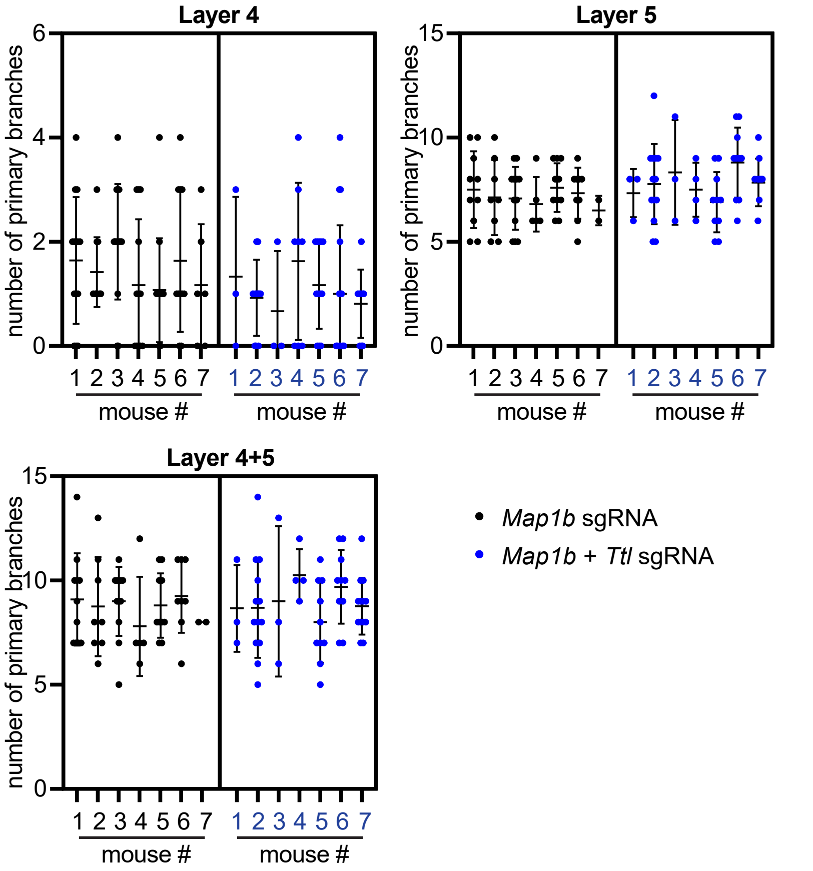
