## Supplementary material for "MAP1B Regulates Cortical Neuron Interstitial Axon Branching Through the Tubulin Tyrosination Cycle": Table EV1

| figures | mouse strain & genotype | plasmids | age of analysis | number of animals | number of neurons | number of neurons per animal | Statistics |
|---|---|---|---|---|---|---|---|
| Figure 1B | WT CD1 | control: pCAG-CreERT2, pEF1-Flex-FlpO, pCAG-FSF-mCherry, pCAG-FSF-GFP | P14 | 4 | 66 | 8-25 neurons/animal | L4 p=0.0124, L5 p=0.0057, L4+5 p=0.0128 |
| | WT CD1 | experimental: pCAG-CreERT2, pEF1-Flex-FlpO, pCAG-FSF-mCherry, pCAG-FSF-GSK3BCA-IRES-GFP | P14 | 8 | 63 | 1-18 neurons/animal | nested t-test (mixed model) |
| Figure 1C | WT CD1 | control: pCAG-CreERT2, pEF1-Flex-FlpO, pCAG-FSF-mCherry, pCAG-FSF-GFP | P14 | 6 | 74 | 10-14 neurons/animal | L4, p=0.9754 |
| | WT CD1 | experimental: pCAG-CreERT2, pEF1-Flex-FlpO, pCAG-FSF-mCherry, pCAG-FSF-GSK3BDN-IRES-GFP | P14 | 6 | 64 | 5-13 neurons/animal | nested t-test (mixed model) |
| Figure 1D, E | *GSK3 α+/-;βfl/+ (control 'het'), GSK3α+/-;βfl/fl and GSK3α+/+;βfl/fl ('Gsk3B KO'), GSK3α-/-;βfl/fl ('double KO')* | pCAG-CreERT2, pEF1-Flex-FlpO, pCAG-FSF-mCherry | P14+P21 | 22 (4xhet, 11xGsk3B KO, 8x double KO) | 131 (28xhet, 71xGsk3B KO, 25x double KO) | 1-13 neurons/animal | nested ANOVA (mixed model), L4 p=0.5462, L5 p=0.039, L4+L5 p=0.2636 |
| Figure 2A, B | WT CD1 | control: pCAG-GFP, pPrime-dsRed-miR30-shRNA scrambled | P14 | 4 | 29 | 3-19 neurons/animal | L4 p=0.0038 |
| | WT CD1 | experimental: pCAG-GFP, pPrime-dsRed-miR30-shRNA-MAP1B | P14 | 7 | 82 | 4-26 neurons/animal | nested t-test (mixed model) |
| Figure 2C, D | WT CD1 | control: pCAG-CreERT2, pTRE-Flex-FlpO, pCAG-FSF-mCherry-tTA, 2x pX458-sgRNA-control2x-Cas9-T2A-GFP | P14 | 8 | 42 | 1-16 neurons/animal | L4 p<0.0001, L5 p=0.0074, L4+L5 p=0.001 |
| | WT CD1 | experimental: pCAG-CreERT2, pTRE-Flex-FlpO, pCAG-FSF-mCherry-tTA, 2x pX458-sgRNA-MAP1B2x-Cas9-T2A-GFP | P14 | 5 | 76 | 11-21 neurons/animal | nested t-test (mixed model) |
| Figure 2E, F | WT CD1 | control: pCAG-CreERT2, pEF1-Flex-FlpO, pCAG-FSF-mCherry, pCAG-FSF-GFP | P21 | 6 | 27 | 2-8 neurons/animal | L4 p=0.166, L5 p<0.001 |
| | WT CD1 | experimental: pCAG-CreERT2, pEF1-Flex-FlpO, pCAG-FSF-MAP1B-Flag, pCAG-FSF-GFP | P21 | 4 | 25 | 3-8 neurons/animal | nested t-test (mixed model) |
| Figure 3A, B | WT CD1 | control: pCAG-CreERT2, pEF1-Flex-FlpO, pCAG-FSF-MAP1B-Flag, pCAG-FSF-GFP | P14 | 6 | 41 | 1-16 neurons/animal | L4 p=0.0346, L5 p=0.0051, L4+L5 p=0.0033 |
| | WT CD1 | experimental: pCAG-CreERT2, pEF1-Flex-FlpO, pCAG-FSF-MAP1B-dP-Flag, pCAG-FSF-GFP | P14 | 2 | 22 | 7-15 neurons/animal | nested ANOVA (mixed model) with post-hoc Dunnet's test |
| | WT CD1 | experimental: pCAG-CreERT2, pEF1-Flex-FlpO, pCAG-FSF-MAP1B-P-Flag, pCAG-FSF-GFP | P14 | 9 | 91 | 4-13 neurons/animal | additional analysis shown in Fig EV3E: L4 p=0.0184, L5 p=0.2456 |
| Figure 3C, D | WT CD1 | control: pCAG-CreERT2, pEF1-Flex-FlpO, pCAG-FSF-mCherry, pCAG-FSF-GSK3BCA-IRES-GFP | P14 | 6 | 86 | 5-19 neurons/animal | L4 p=0.015, L5 p=0.0024, L4+L5 p=0.0014 |
| | WT CD1 | experimental: pCAG-CreERT2, pEF1-Flex-FlpO, pCAG-FSF-mCherry, pCAG-FSF-GSK3BCA-IRES-GFP, pCAG-FSF-MAP1B-dP-Flag | P14 | 5 | 48 | 4-18 neurons/animal | nested t-test (mixed model) |
| Figure 4B | WT CD1 | control: pCAG-GFP, pMini-TagRFP-T_A1aY1, pX330-2xsgRNARosa26-Cas9 | P4 | 3 | 25 | 2, 9, 14 neurons/animal | L4 vs L5 signal, p=0.0025, Wilcoxon test |
| Figure 4C, D | WT CD1 | control: pCAG-CreERT2, pEF1-Flex-FlpO, pCAG-FSF-mCherry, pCAG-FSF-GFP | P14 | 5 | 42 | 1-16 neurons/animal | L4 p<0.0001, L5 p=0.23, L4+L5 p=0.0074 |
| | WT CD1 | experimental: pCAG-CreERT2, pEF1-Flex-FlpO, pCAG-FSF-mCherry, pCAG-FSF-VASH1-IRES-GFP, pCAG-FSF-SVBP-IRES-GFP | P14 | 6 | 87 | 9-22 neurons/animal | nested ANOVA (mixed model) with post-hoc Dunnet's test |
| | WT CD1 | experimental: pCAG-CreERT2, pEF1-Flex-FlpO, pCAG-FSF-mCherry, pCAG-FSF-TTL-IRES-GFP | P14 | 9 | 87 | 4-18 neurons/animal | |
| Figure 4F, G | WT CD1 | pCAG-CreERT2, pEF1-Flex-FlpO, pCAG-FSF-mCherry, pCAG-FSF-TTL-IRES-GFP | P14 | 5 | 33 | 5-14 neurons/animal | L4, p=0.0343 |
| | WT CD1 | pCAG-CreERT2, pEF1-Flex-FlpO, pCAG-FSF-mCherry, pCAG-FSF-TTL-DN-IRES-GFP | P14 | 3 | 21 | 2-17 neurons/animal | nested t-test (mixed model) |
| Figure 5A, B | WT CD1 | control: pCAG-CreERT2, pTRE-Flex-FlpO, pCAG-FSF-mCherry-tTA, 2x pX458-sgRNA-control2x-Cas9-T2A-GFP | P14 | 6 | 15 | 1-5 neurons/animal | L4 p=0.042, L5 p=0.882, L4+L5 p=0.723 |
| | WT CD1 | experimental: pCAG-CreERT2, pTRE-Flex-FlpO, pCAG-FSF-mCherry-tTA, 2x pX458-sgRNA-TTL2x-Cas9-T2A-GFP | P14 | 8 | 88 | 3-23 neurons/animal | nested ANOVA (mixed model) with post-hoc Dunnet's test |
| | WT CD1 | experimental: pCAG-CreERT2, pTRE-Flex-FlpO, pCAG-FSF-mCherry-tTA, 2x pX458-sgRNA-SVBP2x-Cas9-T2A-GFP | P14 | 7 | 55 | 3-23 neurons/animal | |
| Figure 5C, D | WT CD1 | control: pCAG-CreERT2, pEF1-Flex-FlpO, pCAG-FSF-mCherry, pCAG-FSF-GSK3BCA-IRES-GFP, pX458-sgRNA control-Cas9-T2A-GFP | P14 | 6 | 20 | 1-6 neurons/animal | L4 p=0.0123, L5 p=0.0942, L4+L5 p=0.0021 |
| | WT CD1 | experimental: pCAG-CreERT2, pEF1-Flex-FlpO, pCAG-FSF-mCherry, pCAG-FSF-GSK3BCA-IRES-GFP, pX458-sgRNA-TTL-Cas9-T2A-GFP | P14 | 7 | 46 | 1-12 neurons/animal | nested t-test (mixed model) |
| Figure 6A, B | WT CD1 | control: pCAG-CreERT2, pTRE-Flex-FlpO, pCAG-FSF-mCherry-tTA, 2x pX458-sgRNA-MAP1B2x-Cas9-T2A-GFP | P14 | 7 | 83 | 2-12 neurons/animal | L4 p=0.0319, L5 p=0.081, L4+L5 p=0.88 |
| | WT CD1 | experimental: pCAG-CreERT2, pTRE-Flex-FlpO, pCAG-FSF-mCherry-tTA, 2x pX458-sgRNA-MAP1B2x-Cas9-T2A-GFP, pX458-sgRNA-TTL2x-Cas9-T2A-GFP | P14 | 7 | 72 | 3-16 neurons/animal | nested t-test (mixed model) |
| Figure EV1B | WT CD1 | control: pCAG-CreERT2, pEF1-Flex-FlpO, pCAG-FSF-mCherry, pCAG-FSF-GFP | P14 | 6 | 65 | 4-15 neurons/animal | L4, p=0.0022 |
| | WT CD1 | experimental: pCAG-CreERT2, pEF1-Flex-FlpO, pCAG-FSF-mCherry, pCAG-FSF-GSK3BCA(human)-IRES-GFP | P14 | 10 | 100 | 2-24 neurons/animal | nested t-test (mixed model) |
| Figure EV1C | WT CD1 | control: pCAG-CreERT2, pEF1-Flex-FlpO, pCAG-FSF-mCherry, pCAG-FSF-GFP | P14 | same as Figure S1B | | | n/a |
| | WT CD1 | experimental: pCAG-CreERT2, pEF1-Flex-FlpO, pCAG-FSF-mCherry, pCAG-FSF-GSK3BCA-IRES-GFP | P14 | same as Figure S1B | | | n/a |
| Figure EV1D | WT CD1 | control: pCAG-CreERT2, pEF1-Flex-FlpO, pCAG-FSF-mCherry, pCAG-FSF-GFP | P56 | 1 | 4 | 4 neurons/animal | L4 p=0.2927, L5 p=0.2996, L4+5 p=0.2431 |
| | WT CD1 | experimental: pCAG-CreERT2, pEF1-Flex-FlpO, pCAG-FSF-mCherry, pCAG-FSF-GSK3BCA-IRES-GFP | P56 | 3 | 10 | 3-4 neurons/animal | nested t-test (mixed model) |
| Figure EV1F | *GSK3 α-/-;βfl/+* | pCAG-CreERT2, pEF1-Flex-FlpO, pCAG-FSF-mCherry | P14+P21 | 2 | 19 | 7-12 neurons/animal | nested t-test, L4 p=0.622, L5 p=0.56, L4+5 p=0.717 |
| Figure EV1G, H | *GSK3α+/-;βfl/fl and GSK3α+/+;βfl/fl ('Gsk3B KO')* | pCAG-CreERT2, pEF1-Flex-FlpO, pCAG-FSF-mCherry | P14+P21 | 3 | 9 | 2-4 neurons/animal | neurons are from the figure 1D, p = 0.03 (number of dendritic intersections at the level of AIS) |
| | *GSK3α-/-;βfl/fl ('double KO')* | pCAG-CreERT2, pEF1-Flex-FlpO, pCAG-FSF-mCherry | P14+P21 | 4 | 12 | 2-5 neurons/animal | nested t-test (mixed model) |
| Figure EV2C | *βcatenin fl/+ or APC fl/+* | pCAG-CreERT2, pEF1-Flex-FlpO, pCAG-FSF-mCherry | P14 | at least 3 from each group | n/a | | |
| Figure EV2D | WT CD1 | control: pCAG-GFP, pPrime-dsRed-miR30-shRNA scrambled | P14 | at least 3 from each group | n/a | | |
| | WT CD1 | experimental: pCAG-GFP, pPrime-dsRed-miR30-shRNA-Macf1 or Clasp1 or Clasp2 | P14 | at least 3 from each group | n/a | | |
| Figure EV3B | WT CD1 | pCAG-mCherry, pMini-donor, pX330-2xsgRNA-Cas9 | P14 | at least 3 from each group | n/a | | |
| Figure EV3C, D | WT CD1 | pCAG-mCherry, pMini-donor, pX330-2xsgRNA-Cas9 | P4, P14 | at least 3 from each group | n/a | | |
| Figure EV3E | WT CD1 | comparison of data from Fig. 3A & Fig. 2E | P14+P21 | | | | L4 p=0.0184, L5 p=0.2456, nested t-test (mixed model) |
| Figure EV4C, D | WT CD1 | pCAG-CreERT2, pEF1-Flex-FlpO, pCAG-FSF-mCherry, pCAG-FSF-TTL-IRES-GFP or pCAG-FSF-Vash-IRES-GFP+pCAG-FSF-SVBP-IRES-GFP | P14 (samples from Fig. 4C) | | | ctrl 10&12 neurons, SVBP 4 neurons, TTL 3 neurons | ctrl vs SVBP p=0.0093, ctrl vs TTL p=0.0225, unpaired t-test |
| Figure EV4E, F | WT CD1 | 2x pX458-sgRNA-TTL2x-Cas9-T2A-GFP | P7 | 3 | 18 control, 21 exper | 6-13 neurons/animal | ctrl (GFP-) vs TTL (GFP+) p<0.001 unpaired t-test |
| | WT CD1 | 2x pX458-sgRNA-SVBP2x-Cas9-T2A-GFP | P7 | 3 | 21 control, 22 exper | 3-13 neurons/animal | ctrl (GFP-) vs TTL (GFP+) p<0.001 unpaired t-test |
| Figure EV5A, B | WT CD1 | control: pCAG-CreERT2, pTRE-Flex-FlpO, pCAG-FSF-mCherry-tTA, 2x pX458-sgRNA-control2x-Cas9-T2A-GFP | P14 | 4 | 55 | 1-27 neurons/animal | L4 p=0.008, L5 p=0.638, L4+L5 p=0.875 |
| | WT CD1 | experimental: pCAG-CreERT2, pTRE-Flex-FlpO, pCAG-FSF-mCherry-tTA, pX458-sgRNA-SVBP-Cas9-T2A-GFP, pX458-sgRNA-MATCAP-Cas9-T2A-GFP | P14 | 6 | 109 | 4-48 neurons/animal | nested t-test (mixed model) |
| Figure EV5C | WT CD1 | comparison of data from Fig. 5A & Suppl. Fig. 5A | P14 | | | | L4 p=0.229, nested t-test (mixed model) |
| | | TOTAL | | 233 | 2003 | | |

**POST-HOC TEST DETAILS**

**For Figure 1E:**

| Dunnett's multiple comparisons test | Mean Diff. | 95.00% CI of diff. | Below threshold? | Summary | Adjusted P Value |
|---|---|---|---|---|---|
| control-L4 vs. GSK3BKO-L4 | 0.2023 | -0.4984 to 0.9031 | No | ns | 0.7111 |
| control-L4 vs. doubleKO-L4 | -0.09443 | -0.9068 to 0.7180 | No | ns | 0.9434 |
| control-L5 vs. GSK3BKO-L5 | 0.4671 | -1.074 to 2.008 | No | ns | 0.6778 |
| control-L5 vs. doubleKO-L5 | 1.717 | 0.07202 to 3.361 | Yes | * | **0.0405** |
| control-L4+L5 vs. GSK3BKO-L4+L5 | 0.4956 | -1.404 to 2.396 | No | ns | 0.7443 |
| control-L4+L5 vs. doubleKO-L4+L5 | 1.378 | -0.6691 to 3.425 | No | ns | 0.2071 |

**For Figure 3B:**

| Dunnett's multiple comparisons test | Mean Diff. | 95.00% CI of diff. | Below threshold? | Summary | Adjusted P Value |
|---|---|---|---|---|---|
| L4-MAP1B vs. L4-MAP1B-dP | -0.2542 | -0.8936 to 0.3851 | No | ns | 0.5353 |
| L4-MAP1B vs. L4-MAP1B-P | -0.5461 | -1.008 to -0.08380 | Yes | * | **0.0213** |
| L5-MAP1B vs. L5-MAP1B-dP | -0.348 | -1.537 to 0.8410 | No | ns | 0.7116 |
| L5-MAP1B vs. L5-MAP1B-P | -1.183 | -1.937 to -0.4298 | Yes | ** | **0.0032** |
| L4+L5-MAP1B vs. L4+L5-MAP1B-dF | -1.114 | -2.577 to 0.3491 | No | ns | 0.1455 |
| L4+L5-MAP1B vs. L4+L5-MAP1B-P | -1.587 | -2.525 to -0.6495 | Yes | ** | **0.0018** |

**For Figure 4D:**

| Dunnett's multiple comparisons test | Mean Diff. | 95.00% CI of diff. | Below threshold? | Summary | Adjusted P Value |
|---|---|---|---|---|---|
| ctrl-4 vs. VASH-L4 | 0.1125 | -0.3031 to 0.5281 | No | ns | 0.75 |
| ctrl-4 vs. TTL-L4 | -0.6232 | -1.039 to -0.2076 | Yes | ** | **0.0021** |
| ctrl-5 vs. VASH-L5 | -0.04149 | -0.9702 to 0.8872 | No | ns | 0.9905 |
| ctrl-5 vs. TTL-L5 | -0.5447 | -1.435 to 0.3460 | No | ns | 0.2569 |
| ctrl-L4+L5 vs. VASH-L4+L5 | 0.262 | -0.7828 to 1.307 | No | ns | 0.7561 |
| ctrl-L4+L5 vs. TTL-L4+L5 | -1.03 | -2.037 to -0.02347 | Yes | * | **0.0449** |

**For Figure 5B:**

| Dunnett's multiple comparisons test | Mean Diff. | 95.00% CI of diff. | Below threshold? | Summary | Adjusted P Value |
|---|---|---|---|---|---|
| control-L4 vs. TTL-sgRNA-L4 | -0.249 | -0.9866 to 0.4886 | No | ns | 0.5854 |
| control-L4 vs. SVBP-sgRNA-L4 | -0.7806 | -1.555 to -0.006097 | Yes | * | **0.0482** |
| control-L5 vs. TTL-sgRNA-L5 | -0.2041 | -1.356 to 0.9473 | No | ns | 0.8629 |
| control-L5 vs. SVBP-sgRNA-L5 | -0.06061 | -1.271 to 1.149 | No | ns | 0.9877 |
| control-L4+5 vs. TTL-sgRNA-L4+5 | -0.4264 | -1.805 to 0.9525 | No | ns | 0.6585 |
| control-L4+5 vs. SVBP-sgRNA-L4+5 | -0.5333 | -1.977 to 0.9104 | No | ns | 0.5663 |
