## Supplementary material for "MAP1B Regulates Cortical Neuron Interstitial Axon Branching Through the Tubulin Tyrosination Cycle": Table EV2

| REAGENT or RESOURCE | SOURCE | IDENTIFIER | |
|---|---|---|---|
| chicken anti-GFP, 1:1000 | AVES | GFP-1010 | |
| Living Colors DsRed Polyclonal antibody, 1:1000 | Clontech | 632496 | |
| rabbit anti-RFP, 1:1000 | Thermo Fisher | R10367 | |
| rat anti-Tubulin-tyrosinated (YL1/2), 1:500 | Millipore | MAB1864 | |
| mouse anti-Flag, 1:200 | Sigma | F1804 | |
| rabbit anti-Flag, 1:200 | Cell signaling | D6W5B | |
| Goat anti-Chicken IgY (H+L), Alexa Fluor 488, 1:1000 | Fisher | A11039 | |
| Goat anti-Rabbit IgG (H+L) Highly Cross-Adsorbed, Alexa Fluor 555, | Fisher | A21429 | |
| Alexa Fluor 647 goat anti-mouse IgG (H+L), 1:1000 | Fisher | A21236 | |
| Alexa Fluor 647 goat anti-rat IgG (H+L), 1:1000 | Fisher | A21247 | |
| DH5α competent bacteria | N/A | N/A | |
| OmniPur® PIPES, Sodium Salt | Millipore Sigma | 6910-OP | |
| Tamoxifen | Sigma | T5648-1G | |
| Corn oil | Sigma | C8267 | |
| DPBS 1x, no calcium, no magnesium | Fisher | 14190250 | |
| Bupivicaine hydrochloride | Sigma | B5274-5G | |
| Buprenorphine ER-LAB 5ml (1mg/ml) | ZooPharm, LLC | N/A | |
| DAPI (4',6-Diamidino-2-Phenylindole, Dihydrochloride) | Fisher | D1306 | |
| FluoroGel with DABCO | Electron microscopy sciences | 17985-01 | |
| REDExtract-N-Amp(TM) | Sigma | R4775-125ML | |
| QuickExtract DNA extraction solution | MRC | DN 131- 25 ml | |
| T4 DNA ligase | NEB | M0202S | |
| T4 DNA ligase buffer | NEB | B0202S | |
| ATP solution | Sigma | A2383-1G | |
| Dinucleotide phosphates | Fisher | 10297018 | |
| NEBuilder® HiFi DNA Assembly Cloning Kit | NEB | E5520S | |
| NucleoBond® Xtra Midi EF | Clontech | 740420.5 | |
| **PLASMIDS** | **SOURCE** | **IDENTIFIER** | |
| pCAG-CreERT2 | Dorskind et al. | | |
| pEF1-Flex-FlpO | Dorskind et al. | | |
| pCAG-FSF-GFP | Dorskind et al. | | |
| pCAG-FSF-mCherry | Dorskind et al. | | |
| pAAV-Tre-Flex-FlpO | Lin et al. 2018 | Addgene no. 118027 | |
| CAG-FSF-RFP-ires-tTA-WPRE | Luo et al. 2016 | Addgene no. 85038 | |
| pCAG-GFP | Dorskind et al. | | |
| pCAG-FSF-GSK3BCA-IRES-GFP human | Dorskind et al. | derived from Addgene no. 14754, Stambolic et al. 1994 | |
| pCAG-FSF-GSK3BCA-IRES-GFP mouse | this paper | | |
| pCAG-FSF-GSK3BDN-IRES-GFP mouse | this paper | | |
| pPrime-dsRed-miR30-shRNA-MAP1B | this paper | | |
| pPrime-dsRed-miR30-shRNA-CLASP1 | this paper | | |
| pPrime-dsRed-miR30-shRNA-CLASP2 | this paper | | |
| pPrime-dsRed-miR30-shRNA-MACF1 | this paper | | |
| pPrime-dsRed-miR30-shRNA-scrambled | this paper | | |
| pX458-sgRNA-control2x-Cas9-T2A-GFP-A | this paper | derived from Addgene no. 48138, Run et al. 2013 | |
| pX458-sgRNA-control2x-Cas9-T2A-GFP-B | this paper | | |
| pX458-sgRNA-control2x-Cas9-T2A-GFP-C | this paper | | |
| pX458-sgRNA-control2x-Cas9-T2A-GFP-D | this paper | | |
| pX458-sgRNA-MAP1B2x-Cas9-T2A-GFP-A | this paper | | |
| pX458-sgRNA-MAP1B2x-Cas9-T2A-GFP-B | this paper | | |
| pCAG-FSF-MAP1B-Flag | this paper | | |
| pCAG-FSF-MAP1B-P-Flag | this paper | | |
| pCAG-FSF-MAP1B-deltaP-Flag | this paper | | |
| pMini-donor-Actin | this paper | | |
| pX330-sgRNA-Actin-Cas9 | this paper | derived from Addgene no. 42230, Cong et al. 2013 | |
| pMini-donor-MAP1B | this paper | derived from pMiniT Vector (NEB, E1202) | |
| pX330-sgRNA-MAP1B-Cas9 | this paper | | |
| pMini-donor-GSK3B | this paper | | |
| pX330-sgRNA-GSK3B-Cas9 | this paper | | |
| pMini-TagRFP-T_A1aY1 | this paper | derived from Addgene no. 158751, Kesarwani et al. 2020 | |
| pX330-2xsgRNA-Rosa26-Cas9 | this paper | | |
| pCAG-FSF-VASH1-IRES-GFP | this paper | | |
| pCAG-FSF-VASH2-IRES-GFP | this paper | | |
| pCAG-FSF-SVBP-IRES-GFP | this paper | | |
| pCAG-FSF-TTL-IRES-GFP | this paper | | |
| pCAG-FSF-TTLDN-IRES-GFP | this paper | | |
| pX458-sgRNA-TTL2x-Cas9-T2A-GFP-A | this paper | | |
| pX458-sgRNA-TTL2x-Cas9-T2A-GFP-B | this paper | | |
| pX458-sgRNA-SVBP2x-Cas9-T2A-GFP-A | this paper | | |
| pX458-sgRNA-SVBP2x-Cas9-T2A-GFP-B | this paper | | |
| pX458-sgRNA-MATCAP2x-Cas9-T2A-GFP | this paper | | |
| **OLIGONUCLEOTIDES** | **NAME** | **SEQUENCE** | **NOTES** |
| genotyping | GSK3A-KO-F | CCC CCA CCA AGT GAT TTC ACT GCT A | Feng-Quan Zhou lab |
| genotyping | GSK3A-KO-R | AAC ATG AAA TTC CGG GCT CCA ACT CT | Feng-Quan Zhou lab |
| genotyping | GSK3B-fl-F | ACA GGC CAC AGG AAG TCA GT | Feng-Quan Zhou lab |
| genotyping | GSK3B-fl-R | TCT GGG CTA TAG CTA TCT AGT AAC | Feng-Quan Zhou lab |
| genotyping | Bcat-fl-F | AAG GTA GAG TGA TGA AAG TTG TT | Jeremy Nathans lab |
| genotyping | Bcat-fl-R | CAC CAT GTC CTC TGT CTA TTC | Jeremy Nathans lab |
| genotyping | APC-fl-F | GTT CTG TAT CAT GGA AAG ATA GGT GGT C | Bart Williams lab |
| genotyping | APC-fl-R | CAC TCA AAA CGC TTT TGA GGG TTG ATT C | Bart Williams lab |
| knockdown | MAP1B-shRNA | GCCCAAGAAAGAAGTGGTTAA | Benoist et al. 2013 |
| knockdown | CLASP1-shRNA | GCCATTATGCCAACTATCTTT | Mimori-Kiyosue et al. 2005 |
| knockdown | CLASP2-shRNA | GTTCAGAAAGCCCTTGATATT | Mimori-Kiyosue et al. 2005 |
| knockdown | MACF1-shRNA | GCAGAGATGTATCATCCATCAA | Ka et al. 2014 |
| CRISPR-knockdown | MAP1B-sgRNA-1 | GGAGCCATCGGGCAGCAT | Bassik Mouse CRISPR Knockout Library, see ref. Morgens et al. 2017 |
| CRISPR-knockdown | MAP1B-sgRNA-2 | GGTTTGTGTCCCACGAT | |
| CRISPR-knockdown | MAP1B-sgRNA-3 | GGAACCCCCAACCTCGGG | |
| CRISPR-knockdown | MAP1B-sgRNA-4 | GTTTCTAAGACGTCACTT | |
| CRISPR-knockdown | control-sgRNA-1 | ggcccccatatatgcca | |
| CRISPR-knockdown | control-sgRNA-2 | aaCCTCAACCATAACCCT | |
| CRISPR-knockdown | control-sgRNA-3 | gccccacccacaataggct | |
| CRISPR-knockdown | control-sgRNA-4 | ATATCCAACTTagccaggcg | |
| CRISPR-knockin | actin-sgRNA-F | AGAAGGCTATAGTCACCTCG | designed in benchling, On- and Off-target score calculated online according to The Doench, Fusi et al. (2016) |
| CRISPR-knockin | actin-sgRNA-R | TCGATCCCCAAGAAAACCCC | |
| CRISPR-knockin | MAP1B-sgRNA-F | ttttttGGGATGTCATTGAA | |
| CRISPR-knockin | MAP1B-sgRNA-R | gggaaagagagtcagcggga | |
| CRISPR-knockin | GSK3B-sgRNA-F | CTGTGTGTTTGAACCCACAT | |
| CRISPR-knockin | GSK3B-sgRNA-R | CACTTTTCAGGAGATCCAAG | |
| CRISPR-knockin | Rosa26-sgRNA-F | cacaagagtagttacttggc | |
| CRISPR-knockin | Rosa26-sgRNA-R | ATATCCAACTTagccaggcg | |
| CRISPR-knockdown | TTL-sgRNA-1 | GGGCATCCTCATCTCCTCAG | Bassik Mouse CRISPR Knockout Library |
| CRISPR-knockdown | TTL-sgRNA-2 | GCCTCTTCCAGTAGCCGG | |
| CRISPR-knockdown | TTL-sgRNA-3 | GGATTGCAAAGTCATCAGC | |
| CRISPR-knockdown | TTL-sgRNA-4 | GCTGGTGAATTACTACCGGG | |
| CRISPR-knockdown | SVBP-sgRNA-1 | GTCTCAGTTTCCTGCACAA | |
| CRISPR-knockdown | SVBP-sgRNA-2 | GCATTCAGAAGCCAAACCA | |
| CRISPR-knockdown | SVBP-sgRNA-3 | TAAAGAACCAGCCTTCAGAG | |
| CRISPR-knockdown | SVBP-sgRNA-4 | GCTCTCAACAGAGTCATGA | |
| CRISPR-knockdown | MATCAP-sgRNA-1 | GGTGCTGGACTCGGGGACAC | Bassik Mouse CRISPR Knockout Library, reference gene name: 4931428F04Rik, see ref. Landskron et al. 2022 |
| CRISPR-knockdown | MATCAP-sgRNA-2 | GCATCCCCGTCCAATCG | |
| CRISPR-knockdown | MATCAP-sgRNA-3 | GCATGTATTTGCGCACGG | |
| CRISPR-knockdown | MATCAP-sgRNA-4 | GAAGGGACTGCCAGAGCCAA | |
| pX plasmid construction | insert1 - F | tggccttttgctggccttttgctcacatgtGAGGGCCTATTTCCCATG | TKIT method reference: Fang et al. 2021 |
| pX plasmid construction | insert1 - R | tagctctaaaacNNNNNNNNNNNNNNNCGGTGTTTCGTCCTTTCCAC | |
| pX plasmid construction | insert2 - F | aggacgaaacaccGNNNNNNNNNNNNNGTTTTAGAGCTAGAAATAGCAAG | |
| pX plasmid construction | insert2 - R | tagctctaaaac**NNNNNNNNNNNNNNN**CGGTGTTTCGTCCTTTCCAC | |
| pX plasmid construction | insert3 - F | aggacgaaacaccG**NNNNNNNNNNNNNNNNNNNNN**GTTTTAGAGCTAG[illegible] | |
| pX plasmid construction | insert3 - R | ccatttaccgtaagttatgtaacgggtaccgatatctagaaaaaagcaccg | |
